# Intranasal administration of SARS-CoV-2 neutralizing human antibody prevents infection in mice

**DOI:** 10.1101/2020.12.08.416677

**Authors:** Hongbing Zhang, Zhiyuan Yang, Jingyi Xiang, Ziyou Cui, Jianying Liu, Cheng Liu

## Abstract

Prevention of SARS-CoV-2 infection at the point of nasal entry is a novel strategy that has the potential to help contain the ongoing pandemic. Using our proprietary technologies, we have engineered a human antibody that recognizes SARS-CoV-2 S1 spike protein with an enhanced affinity for mucin to improve the antibody’s retention in respiratory mucosa. The modified antibody, when administered into mouse nostrils, was shown to block infection in mice that were exposed to high titer SARS-CoV-2 pseudovirus 10 hours after the initial antibody treatment. Our data show that the protection against SARS-CoV-2 infection is effective in both nasal and lung areas 7 days after viral exposure. The modified antibody is stable in a nasal spray formulation and maintains its SARS-CoV-2 neutralizing activity. Nasal spray of the modified antibody can be developed as an affordable and effective prophylactic product to protect people from infection by exposure to SARS-CoV-2 virus in the air.

**One-sentence summary:** A Fc-modified human antibody prevents SARS-CoV-2 viral infection via nasal administration

## Introduction

The number of COVID-19 cases and hospitalizations have surged as increases in social activity and mobility have led to increased incidences of viral transmission. COVID-19 is transmitted primarily from person-to-person through respiratory droplets when an infected person talks, sneezes, or coughs. Infectious droplets can land in the mouths or noses of people who are nearby or possibly be inhaled into the lungs, with the upper respiratory mucosal surfaces being the initial and predominant sites for the viral infection [1, 2]. In addition, airborne transmission of the virus can occur through aerosol particles that linger in the air for longer periods of time and can travel further from their origin than droplets [3, 4]. Face masks are being used as the first line of defense [5–7], but they are passive barriers to infection and their efficacy is imperfect. Therefore, we explored whether directly deploying SARS-CoV-2 neutralizing antibodies into the upper respiratory airway can effectively prevent infection. A nasal spray can then be used to deliver SARS-CoV-2 neutralizing antibodies in a simple and effective manner to provide protection in addition to face masks, and in situations where face mask wearing is impractical.

In humans, IgA is the major antibody isotype secreted in the upper airways; its presence there correlates with resistance to infection by respiratory viruses [8–10]. Antibody delivery to the upper airway mucosal surface, mimicking naturally secreted antibodies, can prevent virus from reaching its target or directly neutralize infectious virus, and may be a useful strategy for prophylaxis. Indeed, antibodies have been shown to provide protection of the respiratory tract from viral infection when given prophylactically [8–10]. While secretory IgA antibodies are more efficient than IgG antibodies in providing effective viral protection, IgG antibodies have a well-established modality for large-scale manufacturing and characterization, both of which are essential for providing an affordable and scalable source of antibodies for a prophylactic approach. IgA antibodies have a unique structure and glycosylation pattern that enables binding to mucins in the airway epithelium, resulting in extension of their half-lives in the mucosa [11]. We hypothesized that if IgG antibodies can be engineered to bind mucin, their half-lives in the mucosa and protection against viral infection might be extended.

Here we show that a SARS-CoV-2 antibody IgG molecule with a genetically engineered Fc domain has significantly enhanced binding affinity to mucin. The modified antibody retains its high binding affinity to its target antigen (S1 spike protein of SARS-CoV-2) and has neutralizing activity against SARS-CoV-2 viral infection. The modified IgG antibody showed improved potency as compared to the unmodified antibody in blocking viral infection in a cell-based assay. More importantly, the modified IgG antibody was shown to protect hACE2-transgenic mice from SARS-CoV-2 pseudovirus infection when administered intranasally. This protection lasted for at least 10 hours even at the lowest antibody concentrations tested.

## Results

A panel of antibody clones was identified from Eureka’s E-ALPHA^®^ phage library through phage display screening for high affinity binding to the Receptor Binding Domain (RBD) of the SARS-CoV-2 S1 spike protein (Fig. 1a). One lead antibody (EU126 clone in human IgG1 format) was identified to have high binding affinity to the S1 spike protein (Fig. 1b, K_D_=0.71 nM). A pseudovirus system with spike-protein-expressing lentivirus and hACE2-expressing cell lines [12, 13] was used to validate the neutralizing activity *in vitro*. The EU126 clone can efficiently inhibit pseudoviral infection of the hACE2-expressing 293F cells (Fig. 1c). Furthermore, EU126 can bind to various mutated forms of the S1 spike protein, as well as inhibit their binding to hACE2. The S1 protein mutations tested include that of the most infectious strain, SARS-CoV-2 D614G (Table 1).

**Figure 1:**
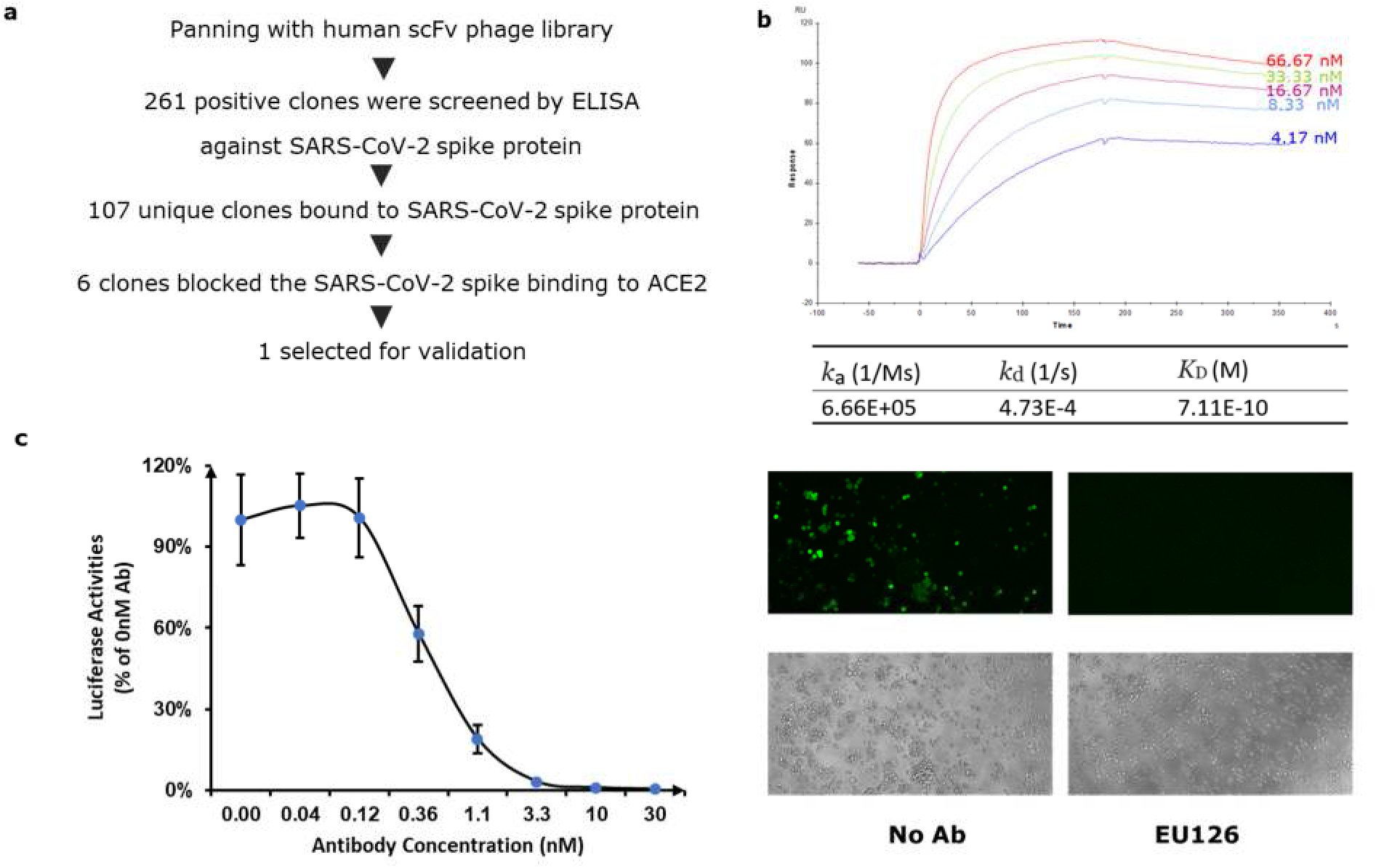
Selection of mAbs that specifically bind SARS CoV-2 S1 and block SARS CoV-2 infection. **1a.** Schematic of the antibody discovery process. **1b.** Multiple cycles of antibody binding kinetics were measured by BiaCore X100 with CAP sensor chip. K_D_ is the binding affinity of antibody. ka, on-rate; k_d_, off-rate; K_D_=k_d_/k_a_. **1c left.** EU126 antibody-mediated neutralization of infection of luciferase-encoding lentiviral particles pseudotyped with spike proteins of SARS-CoV-2. Pseudotyped virus pre-incubated with antibodies at indicated concentrations were used to infect 293F cells overexpressing human ACE2. **1c right.** Pseudotyped virus pre-incubated with no antibody or 10 nM of EU126 antibody were used to infect 293F cells overexpressing human ACE2. The upper panels show GFP fluorescent images of the infected cells; the lower panels show the same cells using light microscopy.

**Table 1.** EU126 binds to various spike proteins or RBD fragments and inhibits their binding to hACE2.

| **SARS-CoV-2 spike variants and other coronavirus spike** | **Binding** | **Inhibition** |
| --- | --- | --- |
| SARS-CoV-2 S1 | + | + |
| SARS-CoV-2 S1 (D614G) | + | + |
| SARS-CoV-2 RBD | + | + |
| SARS-CoV-2 RBD (V367F) | + | + |
| SARS-CoV-2 RBD (N439K) | + | + |
| SARS-CoV-2 RBD (A435S) | + | + |
| SARS-CoV-2 RBD (V483A) | + | + |
| SARS-CoV-2 RBD (K458R) | + | + |
| SARS-CoV-2 RBD (G476S) | + | + |
| SARS-CoV-2 RBD (R408I) | + | + |
| SARS-CoV-2 RBD (V503F) | + | + |
| SARS-CoV-2 RBD (A522V) | + | + |
| SARS-CoV-2 RBD (Y508H) | + | + |
| SARS-CoV-2 RBD (R452R) | + | + |
| SARS-CoV-2 RBD (A520S) | + | + |
| SARS-CoV-2 RBD (I472V) | + | + |
| SARS-CoV-2 RBD (T478I) | + | + |
| SARS-CoV-2 RBD (V341I) | + | + |
| SARS-CoV-2 RBD (F490S) | + | + |
| SARS-CoV-2 RBD (P384L) | + | + |
| SARS-CoV Spike/S1 Protein | - | N/A |
| MERS-CoV Spike/S1 Protein | - | N/A |
| HCoV-HKU1 Spike/S1 Protein | - | N/A |
| HCoV-NL63 Spike/S1 Protein | - | N/A |
| HCoV-229E Spike/S1 Protein | - | N/A |
| HCoV-OC43 Spike Protein | - | N/A |

Mucin glycoproteins produced by mucus-producing cells in the epithelium or submucosal glands are the major macromolecular constituent of mucus [14]. We hypothesized that binding to mucin can potentially extend the retention time of an antibody in the respiratory tract mucosa. We made various modifications of EU126 in the Fc region and tested their binding to mucin. The modifications significantly increased binding to mucin (Fig. 2a).

**Figure 2:**
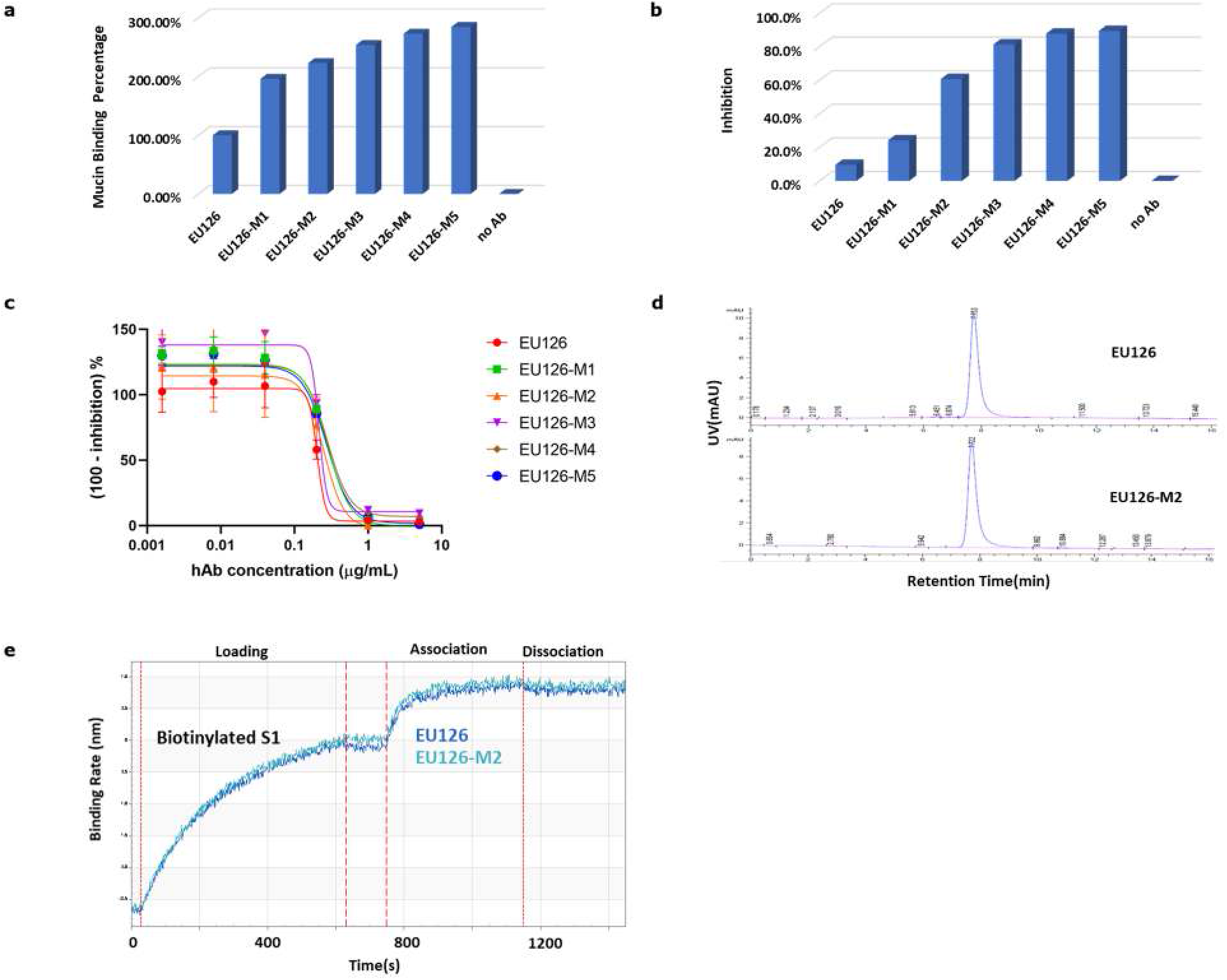
Modifications of EU126 antibody led to significantly increased binding to mucin. **2a.** Modified antibodies have increased binding to mucin as assayed by ELISA. **2b.** Modified antibodies have enhanced neutralization of SARS-CoV-2 infection. Pseudotyped virus were used to infect 293F-ACE2 cells pre-incubated with modified antibodies. Infection of cells was determined by the luciferase level of infected cells using Promega Luciferase Assay. **2c.** The modified antibodies have similar IC50 for S1-hACE2 interaction as EU126. **2d.** Size-exclusion chromatography showed that the M2-modified antibody maintains the same monomeric form as the original clone. **2e.** M2-modified antibody has similar binding affinity to S1 spike protein as EU126 by Bio-layer Interferometry.

Virus neutralization assays are usually performed with antibodies present in solution (Fig. 1c). However, such assays are not suitable for effectively evaluating the antibody binding to mucin-expressing cells in blocking viral infection. An assay was developed to evaluate the effect of direct antibody binding to the cell surface in a SARS-CoV-2 pseudovirus infection (see methods). To test the neutralizing activity of these mucin-bound antibodies, multiple variants of modified antibodies were incubated with cells for 30 minutes to allow binding to the cell surface, the supernatant was removed, and the pseudovirus was added to the antibody-treated cells. As shown in Fig. 2b, all forms of modified EU126 with enhanced mucin-binding affinity showed increased virus neutralization, correlating well with the increased binding affinity to mucin (Fig. 2a). All the modified antibodies have similar IC_50_ for S1-ACE2 interaction as EU126 (Fig. 2c). The modifications in clone EU126-M2 maintain the same monomeric form as the original clone EU126 (Fig. 2d) and has similar binding affinity to the S1 spike protein as EU126 (Fig. 2e). Furthermore, EU126-M2 is stable in a nasal spray formulation for up to 2 weeks at 37°C (data not shown). Therefore, the EU126-M2 antibody was selected for the *in vivo* studies.

In order to test whether antibodies introduced intranasally can provide protection *in vivo,* we established a mouse model for SARS-CoV-2 in which hACE2 transgenic mice [15, 16] were administered intranasally a pseudovirus expressing the SARS-CoV-2 spike protein on the viral surface (Fig. 3a). This pseudovirus harbors a luciferse gene so the infection can be monitored. We tested for virus protection effects with various antibody pretreatment time points, including 24, 10, 6, 4 and 2 hours prior to virus dosing (data not shown). In this model, consistent with the *in vitro* results, EU126-M2 was superior to the parental EU126 antibody in protecting mice from virus infection when antibodies were administered 24 hours prior to virus dosing (Fig. 3b). Thus, the mucin-binding modification in EU126-M2 significantly increased the antibody’s protective effects *in vivo*. A dose titration experiment was performed with different concentrations of EU126-M2 antibody administered into the mouse nostrils before the mice were dosed with the virus intranasally. The untreated infected mice showed a strong luciferase signal in the nasal areas 3 days after virus dosing, while mice treated with 25-200 μg of EU126-M2 had significant reductions in luciferace signals in both nose and lung areas 7 days after virus dosing (Fig. 3c). We tested the duration of protection and found that EU126-M2 provided at least 10 hours of protection (Fig. 3c) against pseudoviral infection. Neither the nasal cavity nor lung areas showed signs of infection in EU126-M2 antibody-treated mice 7 days after virus dosing (Fig. 3c).

**Figure 3:**
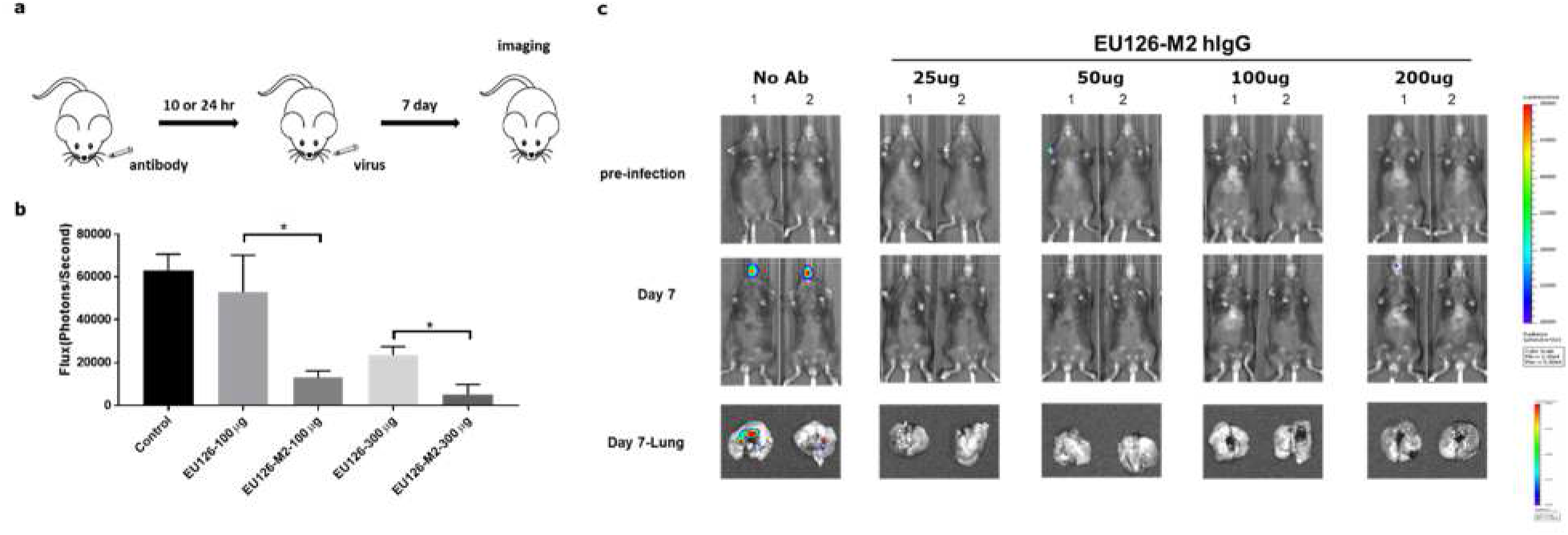
M2-modified EU126 blocks SARS-CoV-2 infection in mouse model. **3a.** Establishment of animal model for SARS-CoV-2 infection. **3b and c.** Comparison of EU126-M2 to EU126 in protection of mice from virus infection. **3b**. EU126-M2 is significantly more effective in inhibiting pseudovirus infection than EU126. Quantification of bioluminescence of SARS-CoV-2 pseudovirus infected mice was carried out at day 7. All data are represented as mean ± S.E. from three biological replicates (*p < 0.05). **3c.** Intranasal administration of 25-200 mg of EU126-M2 blocked pseudovirus infection in both the nostrils and lungs. Antibodies were delivered to mice through nasal administration. 10 hours after antibody delivery, SARS-CoV-2 spike pseudotype lentivirus were delivered to mice through nasal administration. 7 days after nasal administration of virus, bioluminescence imaging was performed for each mouse and its dissected lungs.

## Discussion

Various strategies are being utilized to fight the spread of COVID-19 [17]. Porotto et al. recently reported that an intranasal fusion inhibitory lipopeptide can prevent direct contact transmission of SARS-CoV-2 in ferrets [18]. The dimeric lipopeptide fusion inhibitor works by disrupting the structural rearrangement of the SARS-CoV-2 S1 spike protein that drives membrane fusion of the virus to ACE2-positive cells. The lipo-conjugation of the peptide markedly increased antiviral potency and *in vivo* half-life. The authors proposed to use the lipopeptide as a prophylaxis for SARS-CoV-2 transmission in humans. Other researchers have used a passive immunization approach to harvest immunoglobulin Y antibodies (IgY, the most common immunoglobulin found in birds) from the egg yolks of chickens and formulated them into nasal drops for SARS-CoV-2 prevention [19].

In contrast to these two approaches above, we used a human IgG antibody that binds the RBD domain of the SARS-CoV-2 S1 spike protein. We expect our antibody to block SARS-CoV-2 from binding to ACE2 receptors on cells, entering the cells, and triggering an infection. Despite the differences in modality between a lipopeptide and an antibody, Porotto et al.’s results confirmed that biologically active compounds deposited into the upper respiratory tract can provide effective protection against SARS-CoV-2 infection.

The estimated viral concentrations in a room with a high emitter individual who is coughing frequently can be as high as 7.44×10^6^ copies/m^3^. Regular breathing from a high emitter individual was modeled to result in lower room concentrations of up to 1,248 copies/m^3^ [20].

In this study, we challenged the mice with SARS-CoV-2 infection by directly applying 1×10^7^ pseudovirus particles into the nostrils of mice (Fig. 3c). The inhibition of viral infection by the preadministration of modified IgG antibody against this high viral titer shows the promise of a nasal spray protection even in a worst-case scenario, and could provide a large cushion of protection in most situations of a healthy person encountering an infected individual.

A nasal spray containing IgG antibodies is a complement to vaccines, therapeutics and other preventive measures against the spread of COVID-19. Human IgG antibody nasal sprays have many advantages. The manufacture and scale-up of IgG antibody production is a well-established process [9, 21]. More than two dozen IgG antibodies have been approved by the U.S. Food and Drug Administration (FDA) for human use. The IgG antibody manufacturing process, from cell line development to large scale bioreactor culture, has been optimized to produce high quality and high yield antibodies for use in humans. Multiple clinical trials are currently testing SARS-CoV-2 blocking IgG antibodies as a therapeutic. In November 2020, the FDA granted Eli Lilly and Regeneron emergency use authorization of their IgG antibody therapeutics, for the treatment of confirmed cases of COVID-19.

An IgG in a nasal spray application also has the advantage of a much lower dosage requirement (approximately 10,000 times lower) than an IgG therapeutic which is typically administered intravenously. This will significantly lower the cost and make it affordable for wider use. Other factors that need to be considered include the risk of immunogenicity of the biological compound used in the nasal spray due to the expected long-term repeated use as a prophylaxis. Use of a human IgG antibody significantly reduces the risk of immunogenicity [22, 23]. The long-term stability of the modified IgG antibody at room temperature in the nasal spray formulation makes it easy for daily use and storage.

We believe that a nasal spray containing EU126-M2 human antibodies can be developed as a safe and effective prophylactic product to protect people from infection upon exposure to the SARS-CoV-2 virus, thus slowing the spread of COVID-19.

## Methods

### Phage panning

The E-ALPHA^®^ human scFv antibody phage display libraries (Eureka Therapeutics) were used for the selection of human antibody constructs specific to SARS-CoV-2 spike protein. For protein panning, 5 mg/mL SARS-CoV-2 spike protein (Sino biological, 40591-V08H) was coated on high binding plates (Corning, #3361). Human scFv phage libraries were incubated with a mixture of 6 other coronaviruses for negative selection: SARS-CoV (Sino biological, 40150-V08B1), MERS-CoV (Sino biological, 40069-V08H), HCoV-HKU1 (Sino biological, 40021-V08H), HCoV-NL63 (Sino biological, 40600-V08H), HCoV-229E (Sino biological, 40601-V08H), HCoV-OC43 (Sino biological, 40607-V08B), before adding to SARS-CoV-2 spike protein coated plates. After extended washing with PBS, the bound clones were eluted and used to infect E.coli XL1-Blue cells. The phage clones were expressed in bacteria and purified. Three to four rounds of panning were performed to enrich for scFv phage clones that specifically bound the SARS-CoV-2 spike protein.

### Antibody characterization by ELISA

High binding plates (Corning) were coated with 2 mg/mL coronavirus spike proteins (the spike proteins and their variants are purchased from Sino biological) at least 2 hours at room temperature (RT). Plates were washed with PBS containing 0.05% Tween-20 and blocked with 3% bovine serum albumin in PBS at 4°C overnight. ScFv’s were added to the plates and incubated for 1 hour at RT. Plates were washed three times and incubated with horseradish peroxidase (HRP)-conjugated anti-HA antibody at a 1:2000 dilution. HRP activity was measured at 450 nanometer (nm; OD450) using tetramethylbenzidine (TMB) and an ELISA plate reader. For the inhibition assay, high binding plates (Corning) were coated with 2 mg/mL hACE2-hFc protein in 100 mL PBS for 2 hours at RT. The plates were blocked with 3% BSA overnight at 4°C. Serial dilutions of the antibodies were incubated with SARS-CoV-2-His (2 mg/mL) for 1 hour. The mixture was added to the plates and incubated for 1 hour at RT. Plates were washed 3x with PBST, HRP-Anti-HIS (1:2000) was added and incubated for another hour. HRP activity was measured at 450 nm using TMB and an ELISA plate reader.

### ELISA-based mucin binding assay

In brief, 96-well plates were coated with 50 mg/mL mucin (Sigma, M3895) for 2 hours at RT. The plates were blocked with 3% BSA overnight at 4°C. 5 mg/mL antibodies in 25 mM HEPES (pH6.5) were added to the plates and incubated for 1 hour at RT. After washing with wash buffer (25 mM HEPES, 50 nM NaCl, pH6.5), plates were stained with HRP-conjugated goat anti human IgG and developed using TMB. Absorbance at an optical density of 450 nm was measured.

### Antibody Kinetics by BiaCore

Multiple cycles of antibody binding kinetics were measured by BiaCore X100 with a Sensor Chip CAP. Biotinylated S1 Spike protein was loaded onto the surface of the sensor chip at 5 mg/mL for 90s. Following the loading step, IgG1 antibody was injected onto the Sensor Chip for 180 s at concentrations of 66.67, 33.33, 16.67, 8.33 and 4.17 nM. S1 protein was allowed to associate with the IgG1 antibody for 30 s, and subsequently dissociated for 180 s.

### HLPC-Size exclusion chromatography (SEC)

5 mL of 0.5 mg/mL of antibody was loaded onto a size exclusion column (AdvanceBio SEC Column 300 Å 2.7 μm 4.6 x 300 mm) with a guard column (AdvanceBio SEC Column 300 Å 2.7 μm 4.6 x 50 mm Guard Column) on an Agilent 1260 Infinity HPLC. PBS buffer was passed through the columns at 0.35 mL/min. The UV reading of elution flow was monitored.

### Antibody binding towards S1 Spike Protein by ForteBio Octet

The comparison between EU126 and EU126-M2 binding to spike protein S1 was performed on ForteBio Octet QK (ForteBio) in an 8-channel 96-well plate mode at a shake speed of 1000 rpm. First, the SA sensor tips were dipped into ForteBio kinetics buffer to check the sensor. Then, the sensor tips were exposed to biotinylated spike S1 at 5 mg/mL to saturate SA on the sensor tips. The sensor tips were subsequently dipped into kinetics buffer to elute non-specific binding. Then, the sensor tips with were exposed to the antibodies at 10 mg/mL for association to saturate its binding epitope. Finally, the sensor tips were moved to the buffer for antibody dissociation from antigen.

### Pseudotyped virus neutralization assay

Pseudovirus was generated employing an pCDH lentivirial vector that encodes luciferase and GFP genes pesudotyped with SARS-CoV-2 spike protein. To assess EU126 IgG neutralization, pseudovirus was pre-incubated with varying concentrations of EU126 antibody for 1 hour at room temperature before adding to 293F cells expressing human ACE2. To assess modified EU126 hIgGs, 293F cells expressing human ACE2 were pre-incubated with modified antibodies for 30 min at room temperature followed by removing the supernatant. Pseudovirus was then added to the cells. 48 hours after transduction, infection was determined by detecting luciferase levels in infected cells using Promega Luciferase Assay. The infected 293F cells were imaged by GFP fluorescence.

### Animal study

Female transgenic mice (K18-ACE2) aged 4-6 weeks were used. Antibodies of various concentrations were administered to mice intranasally (20 mL per nostril). 10 hours after antibody administration, SARS-CoV-2 spike pseudotype lentivirus was administered to mice intranasally (20 mL per nostril). 7 days after intranasal administration of virus, bioluminescent imaging was performed for each mouse. After bioluminescence measurement, the lungs were dissected and also imaged.

## Acknowledgement

The authors would like to thank Vivien Chan, Bradley Heller, Sally Fuerstenberg, Mei Hong and Victor Shum for critical review of the manuscript.

## Reference

1. Anderson, E.L., et al., Consideration of the Aerosol Transmission for COVID-19 and Public Health. Risk Anal, 2020. 40(5): p. 902–907.

2. Na, Z., Morphogenesis and cytopathic effect of SARSCoV-2 infection in human airway epithelial cells. 2020.

3. Prather, K.A., C.C. Wang, and R.T. Schooley, Reducing transmission of SARS-CoV-2. Science, 2020. 368(6498): p. 1422–1424.

4. Yuguo, L., Q. hua, and H. Jian, Evidence for probable aerosol transmission of SARS-CoV-2 in a poorly ventilated restaurant. 2020.

5. Stadnytskyi, V., et al., The airborne lifetime of small speech droplets and their potential importance in SARS-CoV-2 transmission. Proc Natl Acad Sci U S A, 2020. 117(22): p. 11875–11877.

6. Chan, J.F., et al., Surgical Mask Partition Reduces the Risk of Noncontact Transmission in a Golden Syrian Hamster Model for Coronavirus Disease 2019 (COVID-19). Clin Infect Dis, 2020. 71(16): p. 2139–2149.

7. Alsved, M. and A. Matamis, Exhaled respiratory particles during singing and talking. 2020. p. 1245–1248.

8. Renegar, K.B., et al., Role of IgA versus IgG in the control of influenza viral infection in the murine respiratory tract. J Immunol, 2004. 173(3): p. 1978–86.

9. Weltzin, R. and T.P. Monath, Intranasal antibody prophylaxis for protection against viral disease. Clin Microbiol Rev, 1999. 12(3): p. 383–93.

10. Ye, J., et al., Intranasal delivery of an IgA monoclonal antibody effective against sublethal H5N1 influenza virus infection in mice. Clin Vaccine Immunol, 2010. 17(9): p. 1363–70.

11. Yue, L. and J. Liang, The Effects of Secretory IgA in the Mucosal Immune System. 2020.

12. Crawford, K.H.D., et al., Protocol and Reagents for Pseudotyping Lentiviral Particles with SARS-CoV-2 Spike Protein for Neutralization Assays. Viruses, 2020. 12(5).

13. Hu, J., et al., Development of cell-based pseudovirus entry assay to identify potential viral entry inhibitors and neutralizing antibodies against SARS-CoV-2. Genes Dis, 2020.

14. Linden, S.K., et al., Mucins in the mucosal barrier to infection. Mucosal Immunol, 2008. 1(3): p. 183–97.

15. McCray, P.B., et al., Lethal infection of K18-hACE2 mice infected with severe acute respiratory syndrome coronavirus. J Virol, 2007. 81(2): p. 813–21.

16. Golden, J.W., et al., Human angiotensin-converting enzyme 2 transgenic mice infected with SARS-CoV-2 develop severe and fatal respiratory disease. JCI Insight, 2020. 5(19).

17. Organization, W.H., COVID-19 STRATEGY UPDATE. 2020.

18. Vries, R.D.d., K.S. Schmitz, and F.T. Bovier, Intranasal fusion inhibitory lipopeptide prevents direct contact SARSCoV-2 transmission in ferrets. 2020.

19. Cohen, J, Can a nose-full of chicken antibodies ward off coronavirus infections? Science: Health, Plants & Animals, Coronavirus Nov. 10, 2020.

20. Riediker, M. and D.H. Tsai, Estimation of Viral Aerosol Emissions From Simulated Individuals With Asymptomatic to Moderate Coronavirus Disease 2019. JAMA Netw Open, 2020. 3(7): p. e2013807.

21. Ruei-Min, L., H. Yu-Chyi, and L. I-Ju, Development of therapeutic antibodies for the treatment of diseases. 2020.

22. Waldmann, H., Human Monoclonal Antibodies: The Benefits of Humanization. Methods Mol Biol, 2019. 1904: p. 1–10.

23. Harding, F.A., et al., The immunogenicity of humanized and fully human antibodies: residual immunogenicity resides in the CDR regions. MAbs, 2010. 2(3): p. 256–65.

